# LabEmbryoCam: An opensource phenotyping system for developing aquatic animals

**DOI:** 10.1101/2023.04.11.536373

**Authors:** Ziad Ibbini, Maria Bruning, Sakina Allili, Luke A Holmes, Ellen Tully, Jamie McCoy, John I. Spicer, Oliver Tills

## Abstract

Phenomics is the acquisition of high-dimensional data on an individual-wide scale and is proving transformational in areas of biological research related to human health including medicine and the crop sciences. However, more broadly, a lack of available transferrable technologies and research approaches is significantly hindering the uptake of phenomics, in contrast to molecular-omics for which transferrable technologies have been a significant enabler. Aquatic embryos are natural models for phenomics, due to their small size, taxonomic diversity, ecological relevance, and high levels of temporal, spatial and functional change. Here, we present LabEmbryoCam, an autonomous phenotyping platform for timelapse imaging of developing aquatic embryos cultured in a multiwell plate format. The LabEmbryoCam capitalises on 3D printing, single board computers, consumer electronics and stepper motor enabled motion. These provide autonomous X, Y and Z motion, a web application streamlined for rapid setup of experiments, user email notifications and a humidification chamber to reduce evaporation over prolonged acquisitions. Downstream analyses are provided, enabling automated embryo segmentation, heartbeat detection, motion tracking, and energy proxy trait (EPT) measurement. LabEmbryoCam is a scalable, and flexible laboratory instrument, that leverages embryos and early life stages to tackle key global challenges including biological sensitivity assessment, toxicological screening and broader engagement with the earliest stages of life.

**Specifications table:** 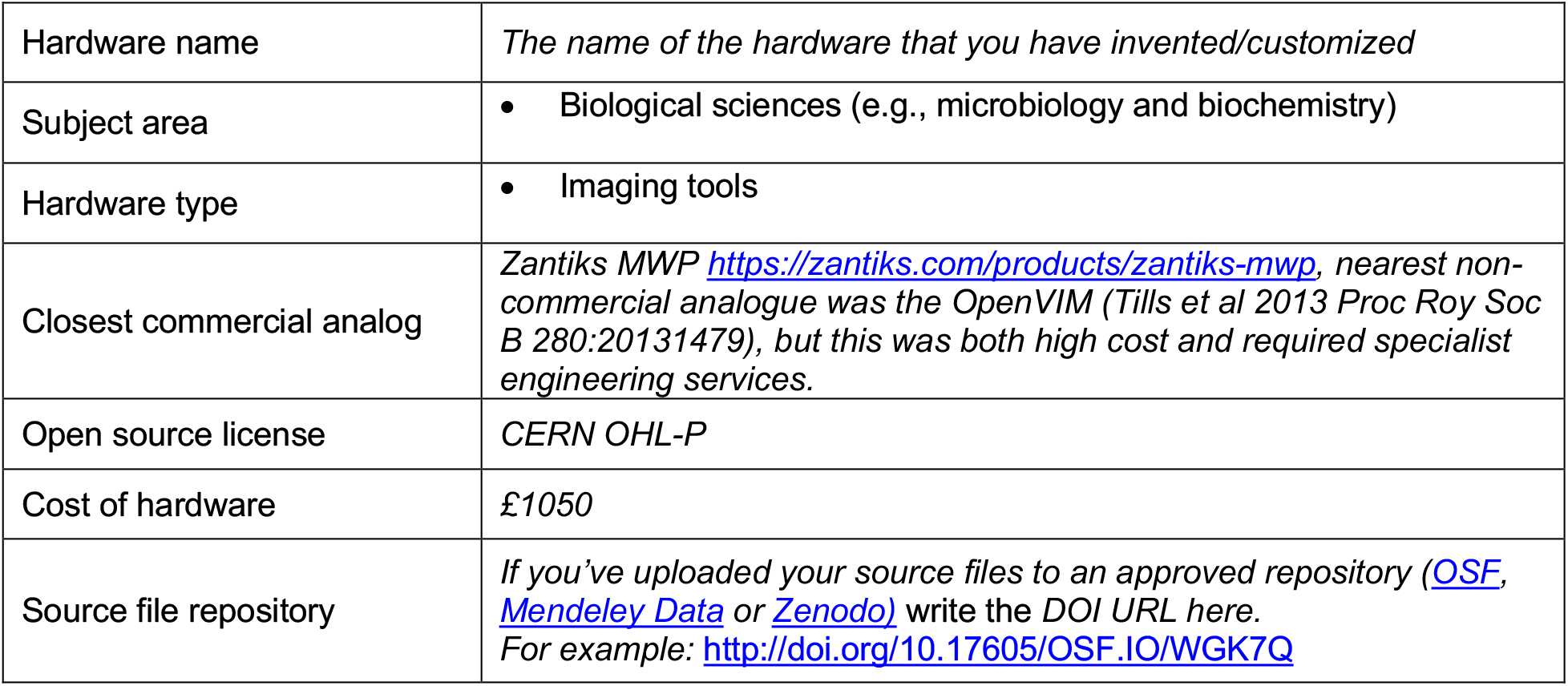

## 1. Hardware in context

LabEmbryoCam is a low-cost phenotyping platform for the automated long term imaging of developing aquatic embryos. The instrument provides a cost-effective solution to the challenge of phenomics – high dimensional phenotyping of organisms [1,2]. A lack of readily available and transferrable tools for phenomics is recognised as a key bottleneck to the advancement of biological research [3]. Aquatic embryos are emerging as natural models for phenomics.They are scalable in both size and taxonomic diversity, and are also the most dynamic stage of life, during which organisms’ typically have heightened sensitivity to their environment [4]. Repeated video observation of individual embryos has proven a powerful method for assessing long term changes in growth and movement, *via* visualization of embryonic development as a timelapse, and as an approach to measuring real-time physiological responses such as heart rate, behaviour and machine proxy traits [5, 6, 7, 8, 9, 10]. The LabEmbryoCam is optimized towards efficiently achieving repeated video image acquisition of embryos in a multiwell plate format for the duration of development, or an experiment. Templates are provided for efficient automated setup of all common multiwell plate formats (6, 24, 48, 96 and 384).

Minimising cost but maximizing versatility were key considerations of both the hardware and software of the LabEmbryoCam, to support its wider adoption at broad numerical and geographical scales. While phenomics is becoming increasingly commonplace in areas of research closely linked to human health, particularly crop sciences and medicine, it remains difficult to integrate more broadly in biological research, except in the case of model species such as zebrafish for which some instruments do exist.

*Daniovision*^©^ is a commercial instrument for tracking small organisms within a multiwell plate, with some support for assisted measurement of heart rate, and Zantiks MWP is another commercial system offering similar functionality. By contrast, the LabEmbryoCam was created to meet a specific research requirement – the high-throughput, long term imaging of developing aquatic embryos, and is both comparatively low cost, and open source. The establishment of transferrable tools and approaches is central to addressing these research requirements, and here, the LabEmbryoCam makes use of the widespread availability of 3D printing, single-board computers (SBCs), low cost electronics and open-source software to offer a solution to the challenge of phenomics of organisms cultured in a multiwell plate format. Unlike molecular-omics, for which one instrument can see application to a broad range of biological systems and research questions, it is well acknowledged that solutions for phenomics are likely to require greater degrees of customization. Opensource hardware and software should therefore be central to successful solutions, and LabEmbryoCam will form a foundation from which variations may emerge to address a range of research challenges, and to capitalize on the availability of novel imaging, motor, sensing, or culture technologies.

## 2. Hardware description

All design files are accessible at *https://doi.org/10.5281/zenodo.7575249*.

The LabEmbryoCam is an imaging instrument built using aluminum extrusion, supported by 3D printed brackets. The frame consists of 2020 (20 × 20 mm cross section) and 2040 (20 × 40 mm cross section) extrusion. Motion is provided using a CoreXY system of moving a carriage – relying on the movement of two stepper motors at the front of the instrument to provide direction in either the X or Y directions. The carriage contains a lens, camera and LED ring light, and also a leadscrew-driven system for Z-travel to provide focus capability. Extensive use is made of 3D printed parts, to improve opportunity for innovation, minimize cost and limit reliance on supply chains. The LabEmbryoCam is highly modular, enabling use of different imaging components, or modification of the design to operate at different scales or resolutions. The instrument tested here uses low-cost optics inverted beneath a multiwell plate, including a Raspberry Pi HQ 12 MP camera, and low cost microscope lens (0.12-1.8 x magnification), with a combined cost of approximately GBP £100.

Electronics are mounted on the rear of the instrument in order to isolate these components from the imaging stage as much as possible. Electronic control systems comprise two Arduino UNO microcontrollers, one for each of the lighting and manual inputs from the joystick and z-control buttons, an XYZ Minitronics microcontroller board, a powered USB hub, an ATX power supply splitter, a Raspberry Pi 4 SBC and an SSD for data storage.

Timelapse imaging of multiwell plate formats can be limited by evaporation altering lighting of embryos, and water chemistry. Consequently, a 3D printed humidification chamber is provided as a cost effective solution to address these limitations and enable longer term autonomous imaging of developing aquatic embryos.

User control of the LabEmbryoCam is achieved through a web application developed in Python using Dash and Flask, among other libraries. The interface is designed to enable efficient setup of long-term image acquisitions, integrating repeated video acquisition of individual embryos over prolonged periods, with autonomous control of the camera, XYZ motion and lighting. Functionality for user updates, *via* email is provided for remote monitoring, and owing to the use of a Dash server based user interface there is potential for accessing instruments in a local area network (LAN) using other devices, as opposed to direct monitor, keyboard and mouse connection.

Key features of the LabEmbryoCam include:

- A significant range of movement 130 mm x 85 mm x 50 mm, with a resolution of 10 microns, highly appropriate to multiwell plate formats.
- Optimised software for efficient acquisition of video at high frame rates and resolutions, using a low powered single board computer (SBC).
- Integration of downstream processes including video compilation and image analysis pipelines, including embryo segmentation, optical flow motion analysis, heart rate quantification and machine proxy traits including energy proxy traits.
- Humidification chamber for stable long-term imaging of organisms cultured in a multiwell plate.
- A capability for stable acquisition of timelapse video over prolonged periods to support downstream analytical processes for producing high-dimensional phenome level datasets.

## 3. Design files

### CAD files

The Autodesk Fusion 360 CAD file for the LabEmbryoCam is available from the paper DOI, and the CAD files are also embedded into the online build guide.

### 3D printing

The LabEmbryoCam incorporates 60 3D printed parts. All STLs are available *via* the Zenodo repository accessible at *https://doi.org/10.5281/zenodo.7575249*, alongside suggested print settings and part orientations.

### Electronics

No custom PCBs are used in this instrument. Figure 12B outlines the connection of the various electronic components to one another, and this is also made clear in both the Python and microcontroller scripts.

### Software and firmware

The LabEmbryoCam is controlled via a web application, built with Python using Dash, running on a Raspberry Pi 4 (SBC), which is in turn connected, *via* USB cables to two Arduino Uno microcontrollers, and a RepRap Minitronics XYZ motion stepper control microcontroller. The firmware for each microcontroller is provided in the Github repository.

A disk image is provided for the Raspberry Pi with all the necessary prerequisite packages preinstalled, however the Github repository for the LabEmbryoCam also includes a requirements.txt file making clear the dependencies.

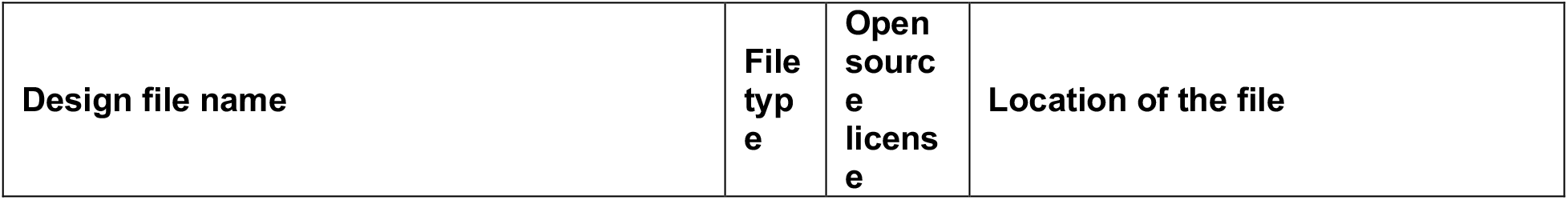
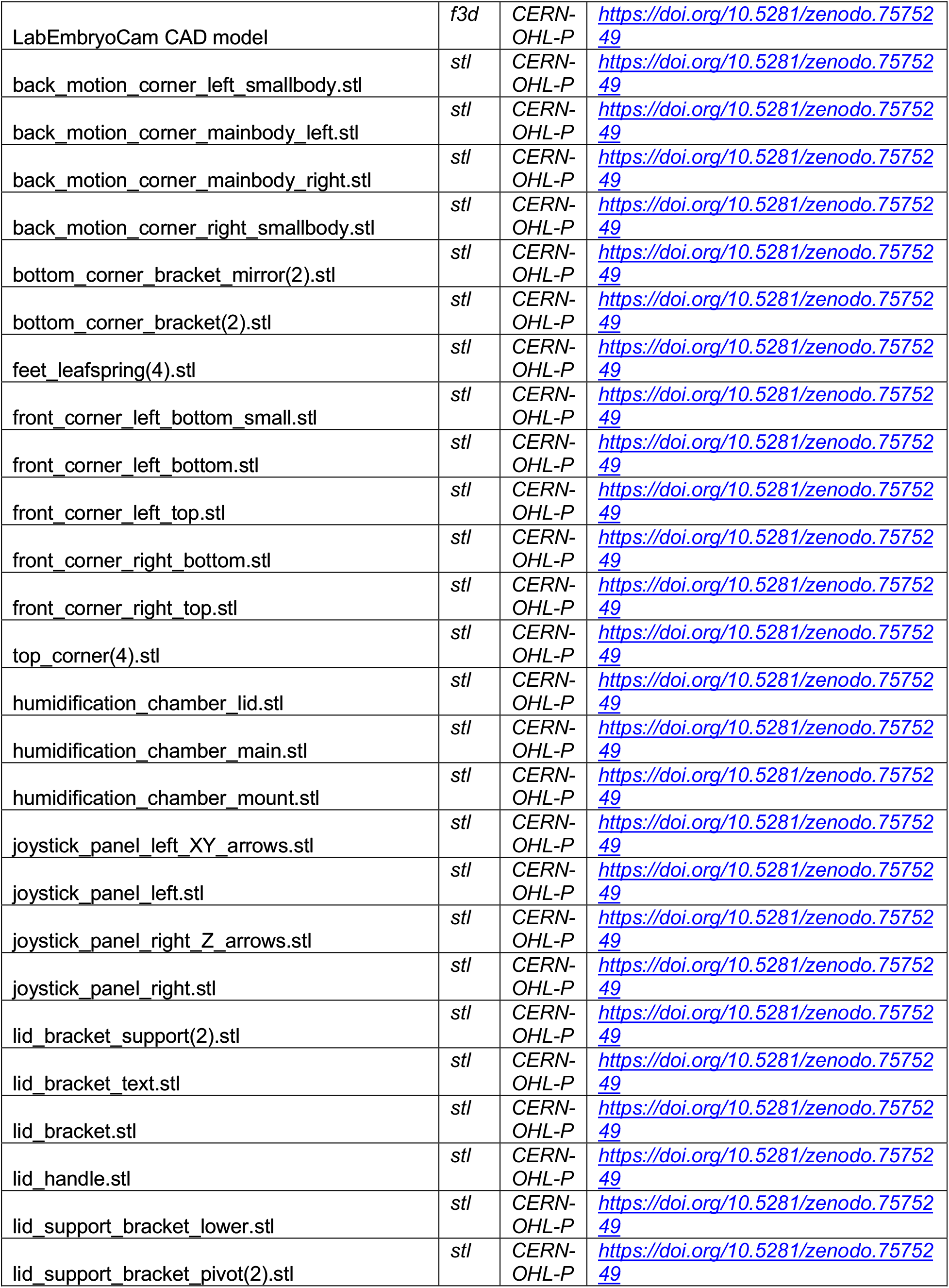
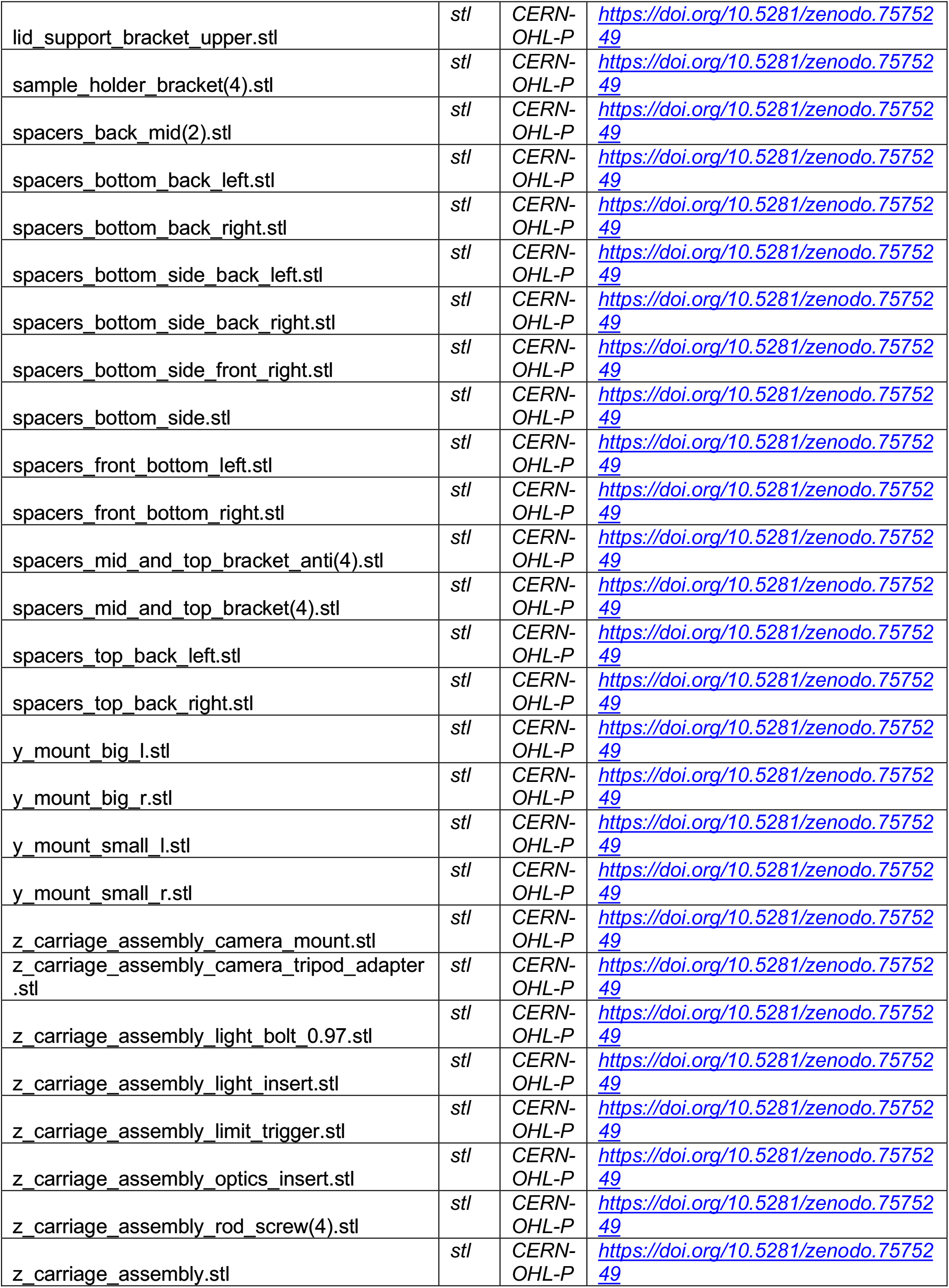
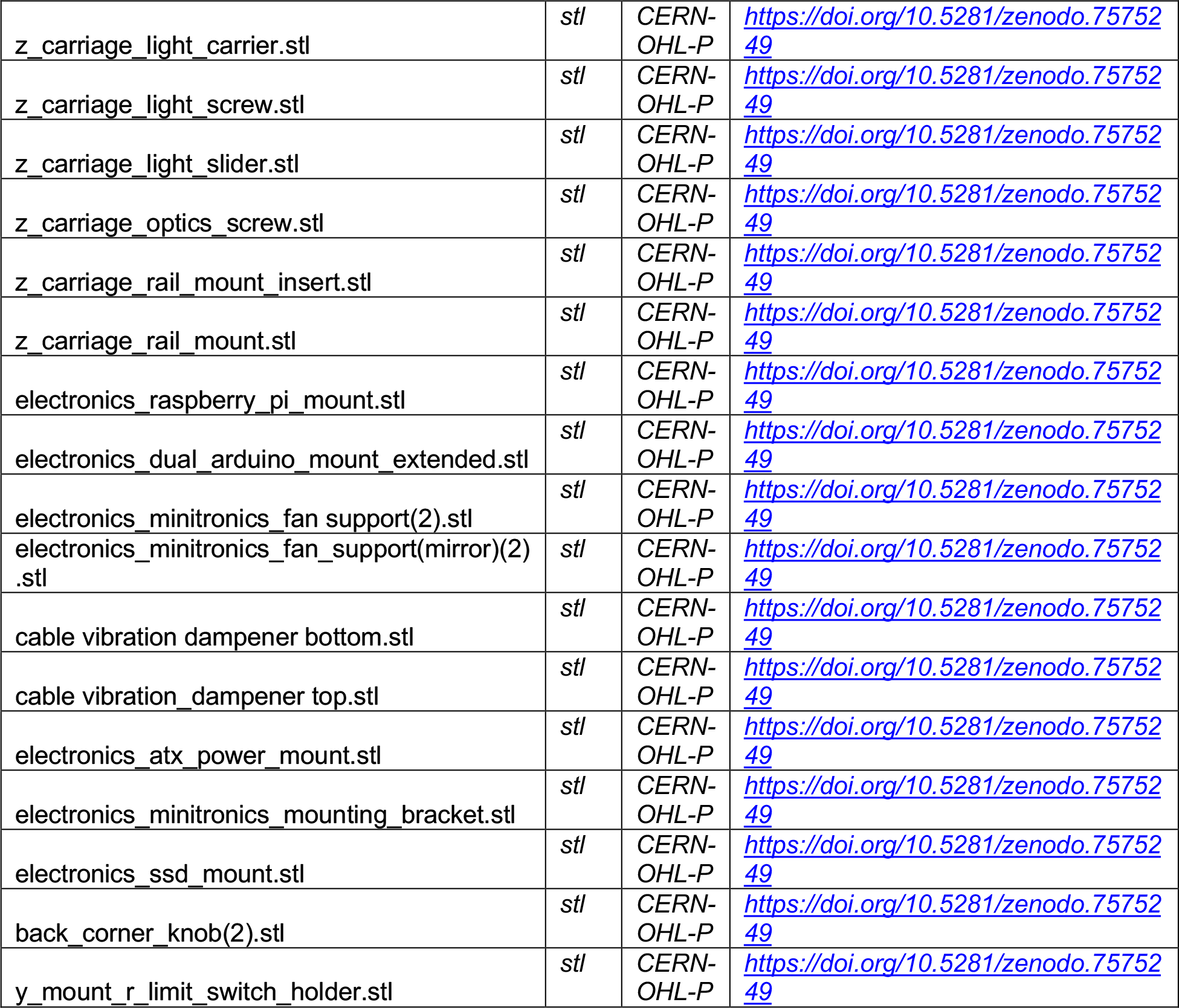

The location of all STLs are detailed in the CAD file, and referred to in the build guide both in this paper and at https://labembryocam.readthedocs.

## 4. Bill of materials

The LabEmbryoCam bill of materials is complex, encompassing > 550 fixings, > 101 3D printed parts and many mechanical and electronic parts, and it is accessible as an *Excel* spreadsheet at https://doi.org/10.5281/zenodo.7575249, providing direct links to product pages, cost, and model numbers. The bill of materials is also accessible available *via* the online build guide. Links in the bill of materials are provided to limit the number of individual orders, and associated shipping charges, but parts will be available from a range of suppliers. Whilst a list of individual fixings is provided and items can be purchased widely, to ease ordering a cost-effective and full collection of fixings for the LabEmbryoCam can be purchased directly at https://www.modelfixings.co.uk/embryophenomics.htm.

The instrument built and tested here is using three widely available MGN12H linear rails and carriages, but alternative higher quality linear rails could also be used, such as those manufactured by Igus.

## 5. Build instructions

The LabEmbryoCam is built using readily-available and low-cost consumer electronics, mechanical parts and 3D printed parts. There is significant scope for changing components, or suppliers due to availability, price, or user requirements.

STLs are provided for all 3D printed parts, but so too are pre-sliced build plates optimized for printing the parts for one LabEmbryoCam. The instrument in this paper was produced using a Prusa MK3S 3D printer and this can construct all of the parts for one LabEmbryoCam in 10 prints, or approximately 500 h of print time. The type of plastic filament used for printing will generally not impact the structure of the main frame. However, chopped carbon-fibre PETG is recommended to improve stiffness and reduce the weight of the parts involved in routing belts (front corners, x-axis, z-carriage and back corners), thereby reducing vibration and increasing accuracy of XYZ movements. Alternatively, to reduce cost, the entire instrument could be produced using PETG, perhaps with higher infill (50 -100%) to maximise rigidity.

The LabEmbryoCam is modelled in Fusion360 and in addition to the model file (*https://doi.org/10.5281/zenodo.7575249*), an online build guide (https://labembryocam.readthedocs) is available with embedded interactive 3D models to visualize the location of individual components.

Upon completion, the LabEmbryoCam is 390 mm deep x 450 mm wide x 475 mm tall. The outer plastic panels and lid are optional, as they are not central to the function of the instrument, but do provide protection to the internal components and help to keep the instrument clean.

The LabEmbryoCam makes use of some ‘home-made’ pulleys, using 3 x M3 washers interleaved between 2 x F623ZZ flanged bearings, as shown below. The belts run over the flanged bearings - so ensure the flanges are on the outer edge to guide the belt (Figure 2A). There are three pulleys in each front corner, two in each X-axis bracket and one in each back corner.

**Figure 1.**
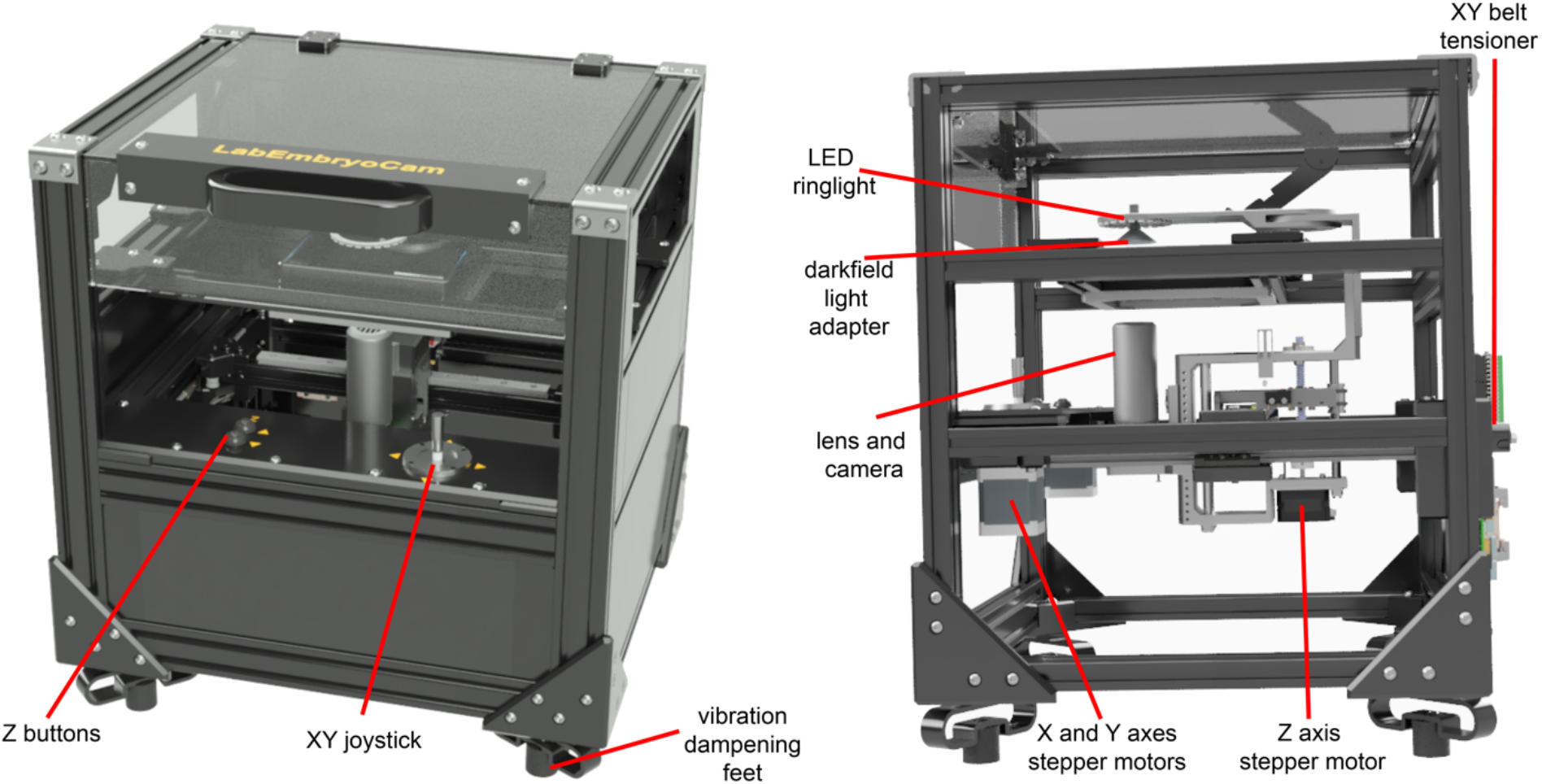
Annotated CAD rendering of an assembled LabEmbryoCam (450 mm wide, 390 mm deep, 475 mm tall) from different orientations. The side image has had the panels and side supports removed to aid visibility of the internal components.

**Figure 2.**
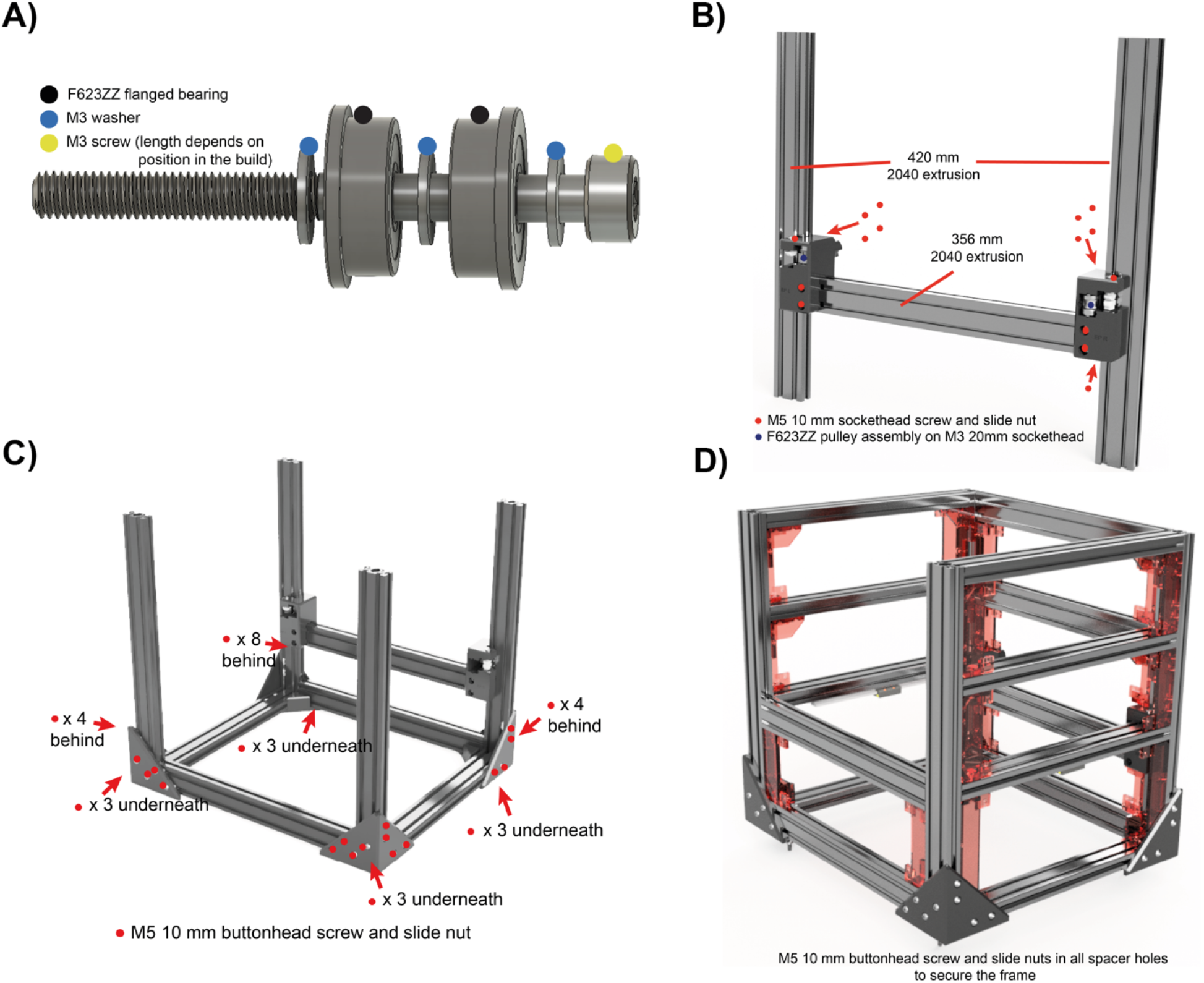
Annotated CAD renders showing different stages of build of the LabEmbryoCam. A) Pulleys built using F623ZZ flanged bearings and M3 washers. These are used for routing the timing belt around the front and back corners, and the X-axis. B) Fixings and pieces of extrusion used at the rear of the instrument for routing and tensioning the belts. C) Fixings used with the upright and bottom pieces of extrusion, secured using the bottom corner 3D printed pieces. D) 3D printed frame spacers and fixings used to attach the remaining pieces of 2040 aluminium extrusion to form the instrument frame.

### Rear corners

See Figure 2B. Begin assembly with the rear cross beam, inserting screws and slide nuts into each of the main back corner 3D printed parts – back-corner-left_mainbody and back-corner-right_mainbody.

Slide these back-corner parts onto each of the two 420 mm 2040 uprights - leaving loose at this stage. Then slide the 356 mm 2040 back lateral into place between the back-corner 3D printed parts. Lay this ‘H’ shape of parts flat and loosely tighten. Don’t worry about the height of the cross bar at this stage - this will be adjusted in subsequent steps.

See Figure 2C. Add nuts and slide bolts to the four corner brackets. Position the four bottom corner brackets flat, taking note of their asymmetry, and slide in the side-to-side and front-to-back pieces of 2040 extrusion. The front-to-back extrusion parts should sit flat on its 40mm side, whereas the side-to-side extrusion sits on its smaller 20mm side. Once the four bottom pieces are inserted the frame should be somewhat stable and square. At this stage, insert the upright pieces of 2040 extrusion. These should reach down to the corner brackets, i.e. the square bottom pieces of extrusion sit against the uprights.

Ensure the parts are fully inserted and square, and then begin tightening up the bolts attached to the slide nuts.

### Frame assembly

See Figure 2D. Before proceeding with assembling the rest of the instrument, it is easiest to loosely attach the two y-axis linear rails with 8 mm countersunk bolts and slide nuts. Tighten just enough to stop them falling off when you turn the extrusion upside down. These rails are what will enable the ‘carriage’ to slide back and forth, providing the y-axis movement. Attach one linear rail to each of the 356 mm 2040 pieces of aluminium extrusion.

Now, insert M5 bolts and slide nuts through the holes of all 3D printed spacer parts before attempting to fit them. Make sure to push the bolts all the way through the 3D printed part and ensure it turns freely before proceeding. Screw the slide nut onto the bolt, but only one turn. Note that the spacers should have the smooth part facing outwards, and that their location is indicated in their naming. If in doubt, consult the CAD model, or figure. The two pieces of extrusion with linear rails should be installed front-to back on the layer above the bottom-most layer. Similarly, if in doubt, check the CAD model. When installing these pieces of extrusion, the linear rail should be on the side of the extrusion closest to the centre of the instrument.

Start assembling the instrument from the bottom-up. First insert the spacers that will sit on the previously assembled bottom square (Figure 2C). Line up the slide nuts in these 3D printed parts with the groove of the extrusion and insert them, before tightening. Note when tightening that the part should be pulled towards the extrusion - i.e. the two are bound together attached. If this does not happen, try untightening the slide nut and repeating the process.

### X-axis

The x-axis relies on a 320 mm piece of 2020 extrusion that moves front to back along the y-axis. It is easiest to build the x-axis outside of the instrument and then attach it to the carriages on the two y-axis linear rails that were previously installed (Figure 3). Furthermore, each end of the X-axis incorporates two pulleys for routing the belts (the pulley assembly is detailed in Figure 2A).

**Figure 3.**
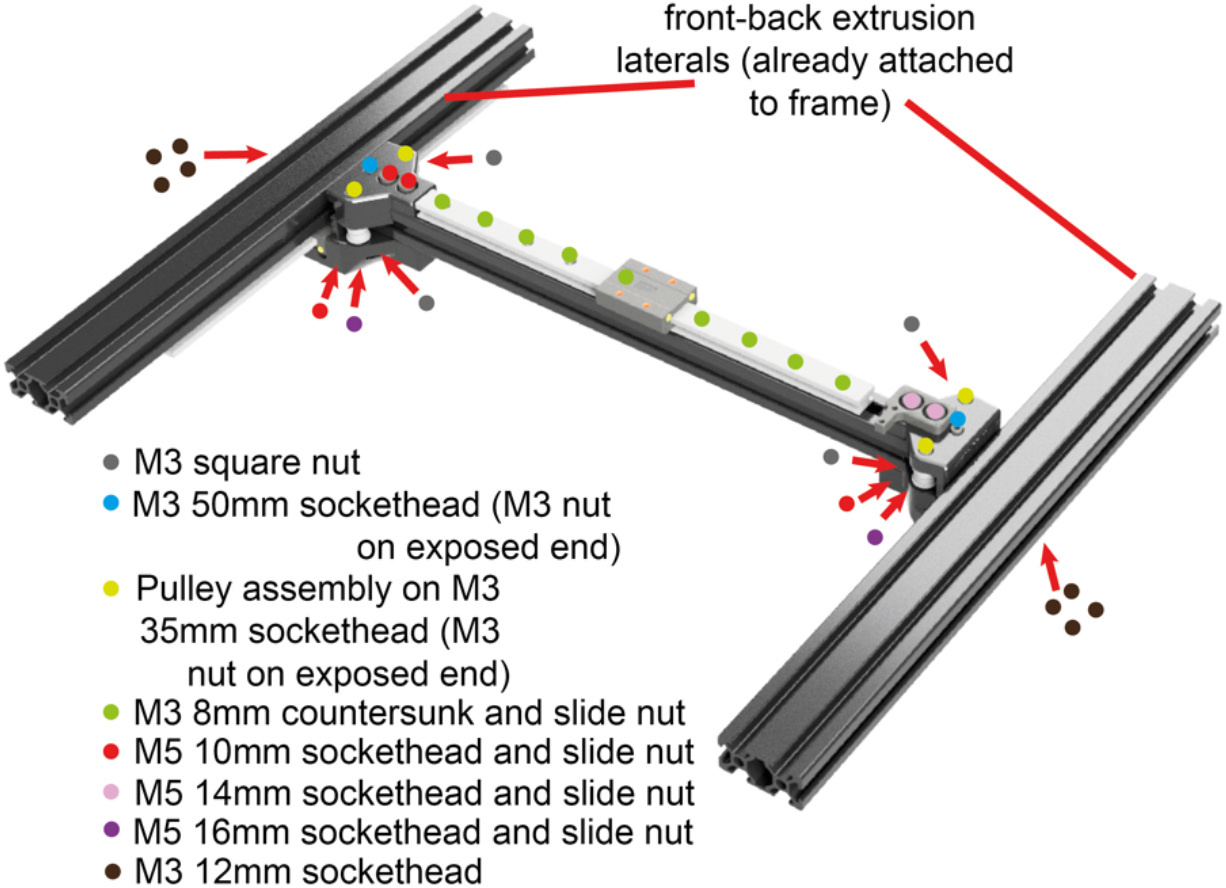
Annotated CAD render of the x-axis assembly. Only the relevant pieces of aluminum extrusion are included in the figure, to make clear the pieces of interest.

### Z-axis

The z-axis incorporates the optics and lighting, but also the mechanism for z-axis motion. This all runs on the z-carriage (Figure 4) from side to side on a linear rail, mounted on a 2020 piece of x-axis extrusion.

The z-axis carriage is the first piece of this assembly to be installed, onto the linear rail carriage. This part stays stationary in the z-direction and provides the base from which the z-assembly then moves up and down. Before attaching the z-carriage part to the linear rail carriage, install cable ties through each of the two holes on both the front and back of the z-carriage. This is harder to do once it is attached to the x-axis linear rail carriage.

### Front corners

The front corners (Figure 7) encompass three pulleys (Figure 2A), a stepper motor and multiple fixings in each corner. The left and right sides are also not symmetrical. The pulleys are responsible for routing the toothed belts, used for transmitting motion from the stepper motors to the X and Y axes.

The lower parts of the front corners should be flush with the top of the lateral 2040 pieces of extrusion to which they are now attached (Figure 7). Begin by attaching these parts to the aluminium extrusion. Note the location of the different screw lengths in the front corners. Insert the M3 35mm screws from underneath until they protrude sufficiently to install the parts required to mount the pulleys (Figure 7).

**Figure 5.**
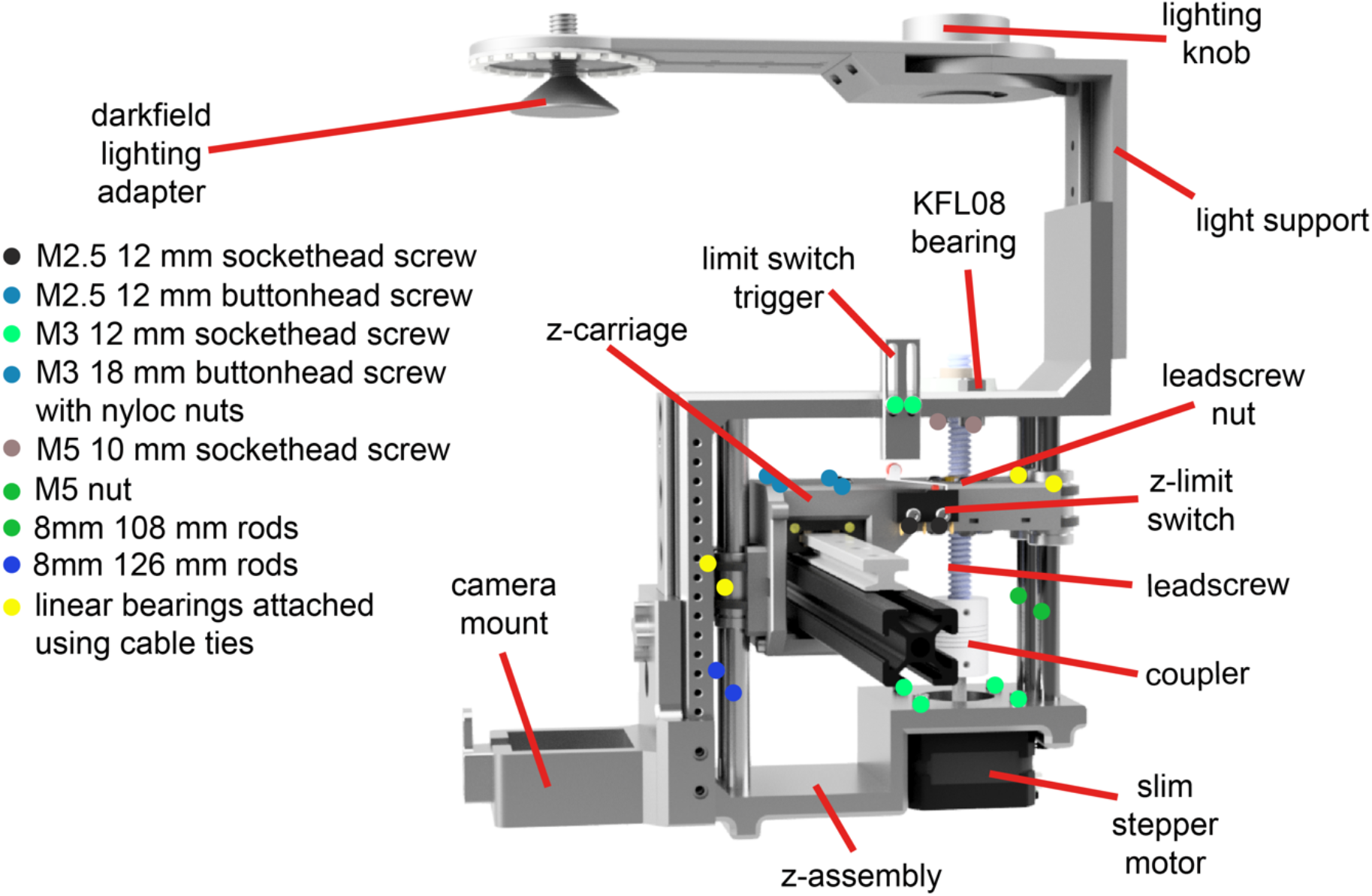
Annotated CAD render of the z-axis assembly. Only the 2020 extrusion and linear rail from the larger assembly are included to aid visibility of the z-axis itself. The sequence of the z-carriage assembly steps is critical to prevent the need for subsequent disassembly at a later stage. 1. Slide the 126 mm smooth rods to the front of the z-axis carriage and insert the linear bearings onto each rod (Figure 6A) 2. Attach the z-axis slimmer stepper motor to the back of the z-axis assembly (Figure 6B). 3. Attach the optics rail to the front of the z-axis assembly (Figure 6C). 4. The two 108 mm rods can now be inserted into the back of the carriage, with a linear bearing on each, but this must be done with the assembly attached to the linear rail (Fig 6D). Cable ties can then be used to secure the linear bearings on both the front and the back rods, to the z-carriage. The leadscrew should then be fed in from the top down through the KFL08 bearing and leadscrew nut to the stepper motor coupler (Figure 5) and secured with the grub screws on the coupler.

**Figure 6.**
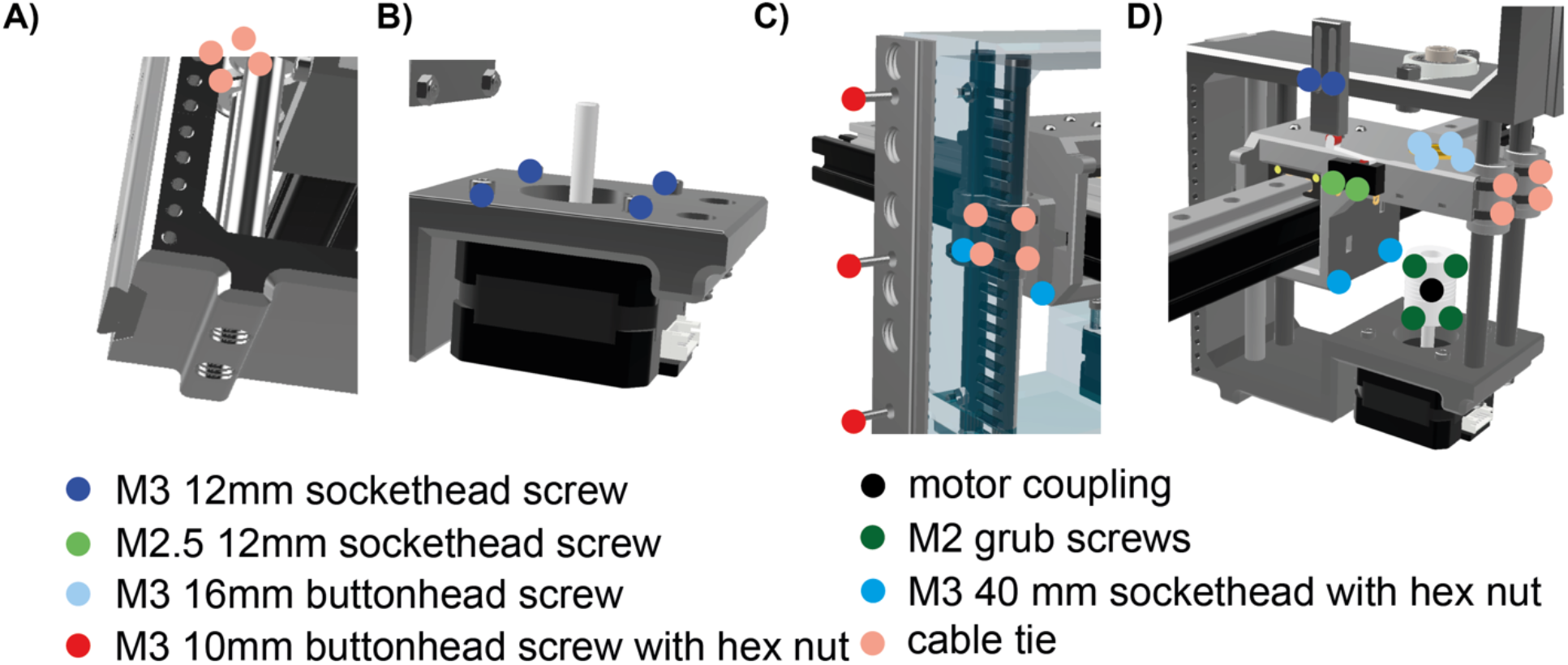
Annotated CAD render of key steps in the process of assembling the z-axis. A) Slide the 126 mm rods into the z-axis assembly and insert the linear bearings onto each rod. B) Attach the slim z-asix stepper motor. C) Attach the optics rail to the front of the z-assembly. D) Insert the two 108 mm rods and attach the linear bearings to the z-carriage with cable ties threaded through the z-carriage.

**Figure 7.**
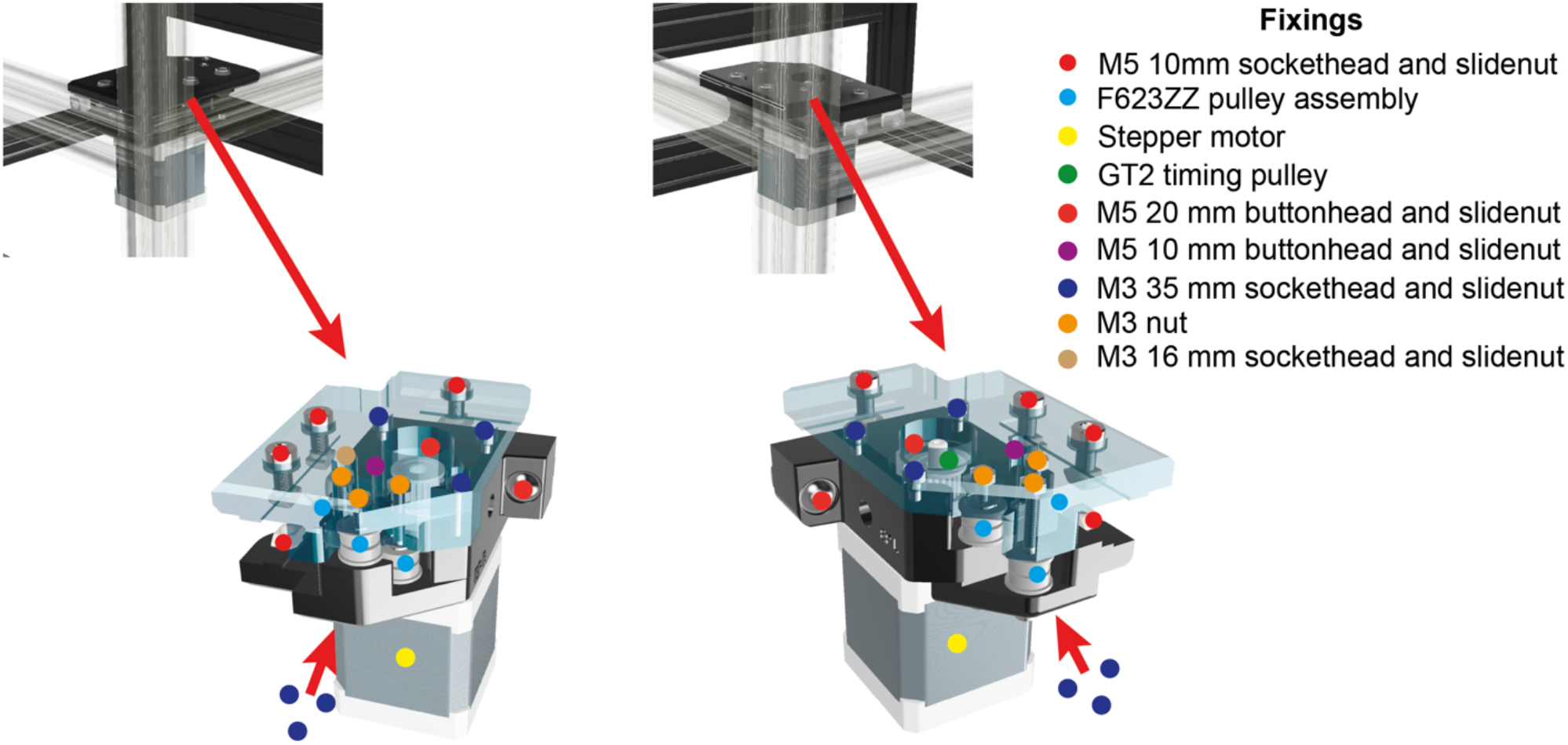
The front corners are responsible for mounting the stepper motors and routing the belts, with an assortment of fixing sizes and types. The top and bottom of the front corner pieces slide together and are subsequently secured with screws. Begin the assembly by attaching the bottom front corner piece on each side of the instrument. Aluminum extrusion is presented as transparent to aid visualization of the location of the front corners.

### Manual controls and vibration insulation

X, Y and Z motion can be controlled from the front panel of the LabEmbryoCam. A toggle switch in the web application activates these controls for user input. Cables must be attached to i) each of the pins on the joystick (2 × 3 cables per axis), and ii) each of the two z buttons (2 × 2 cables) and these pieces of hardware can then be attached to the 3D printed front panels (Figure 8). Ensure that the cables are labelled and then route around the inside of the instrument to the back where they will be attached to one of the two Arduino Uno boards responsible for controlling these manual controls.

**Figure 8.**
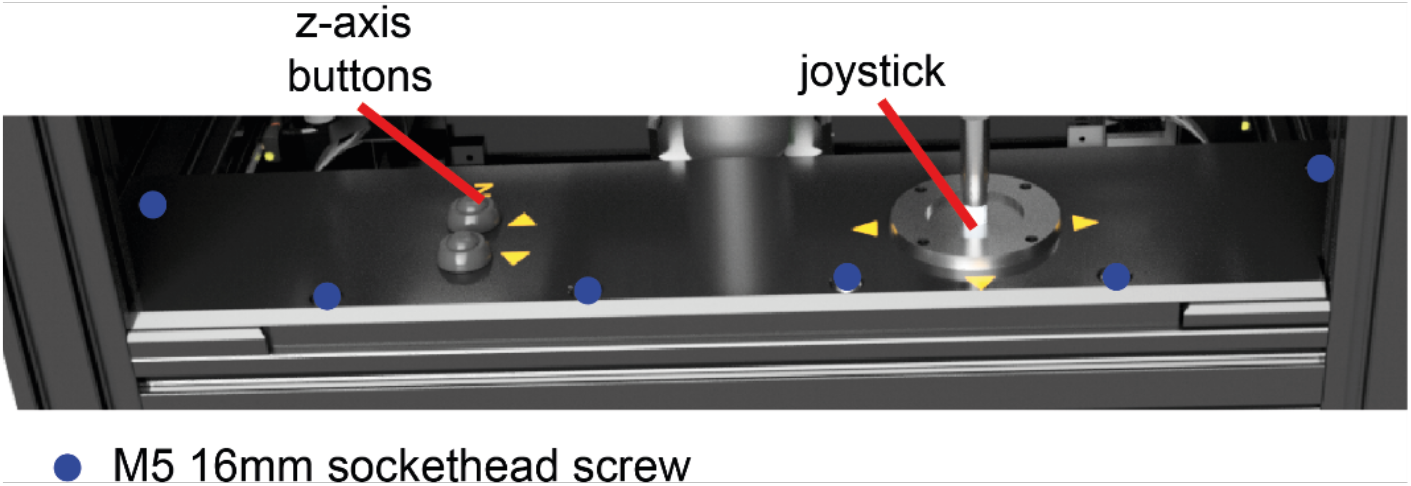
Annotated CAD render of the front panel joystick assembly, mounting the XY joystick, Z buttons and covering the belts at the front of the instrument.

Each bottom corner of the instrument has a vibration insulation foot, consisting of both a Sorbothane^®^ part, and a 3D printed leafspring. These should be attached to the LabEmbryoCam at this stage in the build process (Figure 9).

**Figure 9.**
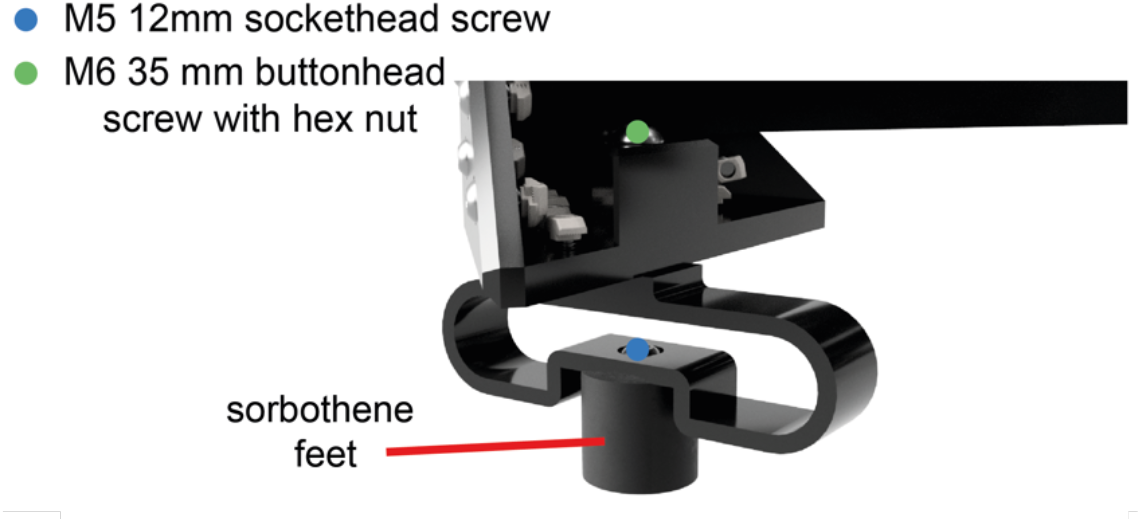
Annotated CAD render of the LabEmbryoCam foot assembly.

### Belt routing

X and Y motion is achieved using a 6mm flexible timing belt, and its correct routing is central to proper function. Two separate belts are attached to the carriage (Figure 10A) and follows a path around both the front and back corners of the LabEmbryoCam.

Insert an M4 40 mm hex screw into the belt tensioners and position loosely in the back corners of the instrument (Figure 2B).

**Figure 10.**
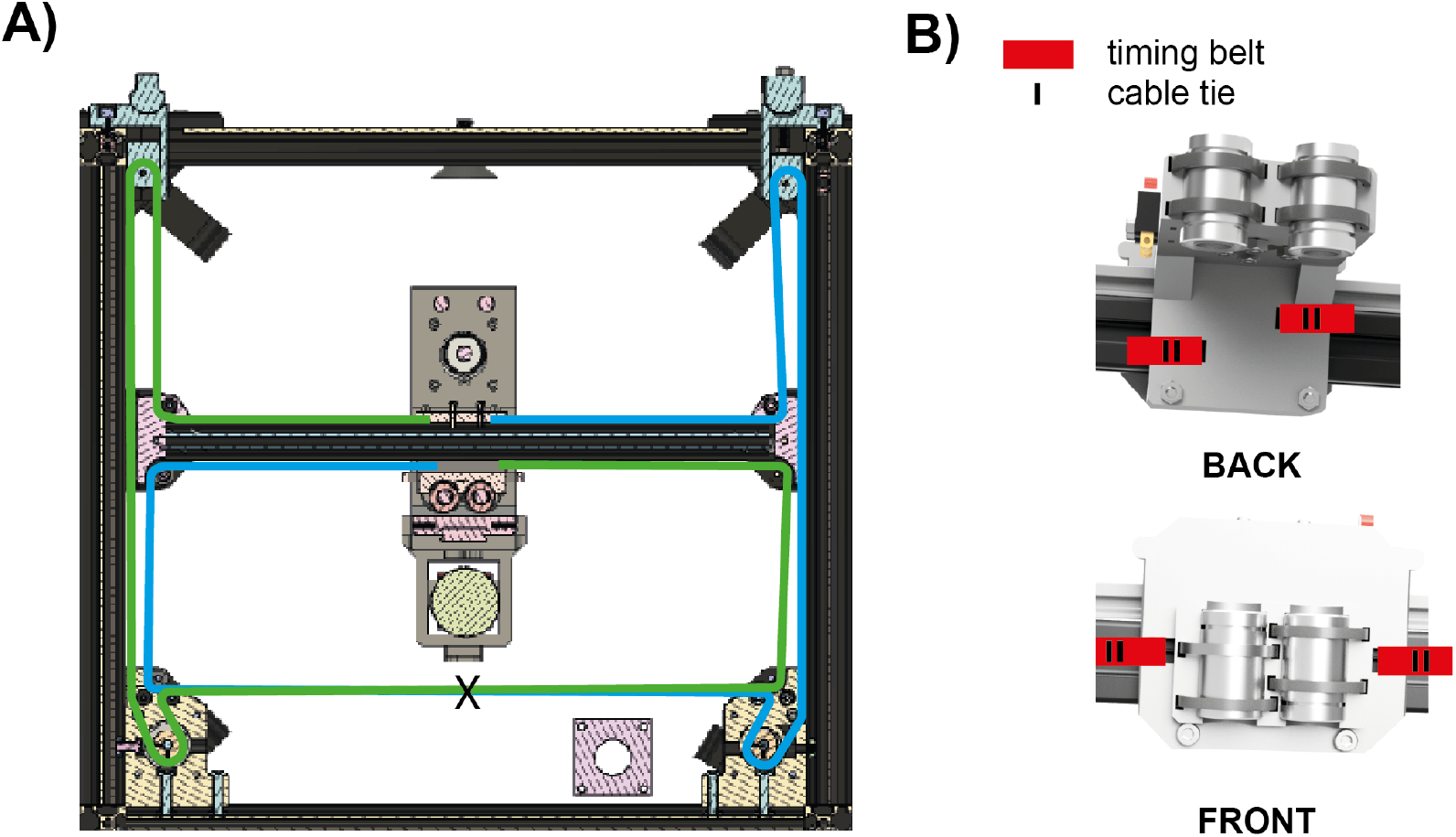
**A)** Cross sectional CAD render of the LabEmbryoCam indicating the routing of the belts in the LabEmbryoCam. Green and blue lines, indicate each of the two belts used to achieve the X and Y movement. ‘X’ indicates where each of the belts should twist by 180? to ensure that the smooth side of the belt is running against the pulleys. **B)** Belt securing locations at the front and back of the z-carriage. The belt should loop around the carriage through the hole indicated, and be secured so that the teeth lock together, using cable ties.

The path of each belt is detailed in Figure 10A, including the location of a 180 degree twist. Each belt both starts and stops at the z-carriage. Here, they should loop behind the carriage, passing through the hole, and be secured with cable ties (see below).

Begin the process of routing by attaching one end of the belt (blue and green in Figure 10A indicate different belts) to the carriage (as seen below). The route for the belts can be seen below. Note that the teeth of the belt must engage with the timing pulleys attached to the stepper motors, but also be rotated by 180 degree at the front of the instrument, to ensure the smooth side is running over the pulleys, rather than the teeth.

The location of where both belts should be rotated is indicated as X in Figure 10A.

Note that each belt loops through the belt tensioners at the back of the instrument. Once the belt has followed the path, loop around the carriage in the same way as for the other end. At this stage, make sure that the belt tensioner (Figure 1) is inserted into the rear of the instrument, and the tensioner knob is screwed on loosely. This is required, to ensure that there is sufficient belt before cutting. Pull the belt tight with a pair of pliers along its route and then secure with a cable tie. Once it is secured with a cable tie to the z-carriage, you can cut the belt. Repeat this process on the other side of the carriage. The belt should be tightened sufficiently that when the stepper motor turns the belt is engaged, but not so much that resistance means that the stepper motors struggle to turn. The tensioner knobs at the back of the instrument, pull the belts backwards and make adjusting belt tension easy.

### Humidification chamber

Evaporation and condensation are common challenges in the maintenance and visualisation of developing aquatic embryos. A cost effective and efficient solution to these challenges is a 3D printed humidification chamber (Figure 11). This humidification module can achieve > 90 % humidity, thereby reducing evaporation to negligible levels. It is not, however, essential to the successful operation of the LabEmbryoCam, for which multiwell plate and Petri dish mounts are available, in addition to a mount for the humidification module described below.

**Figure 11.**
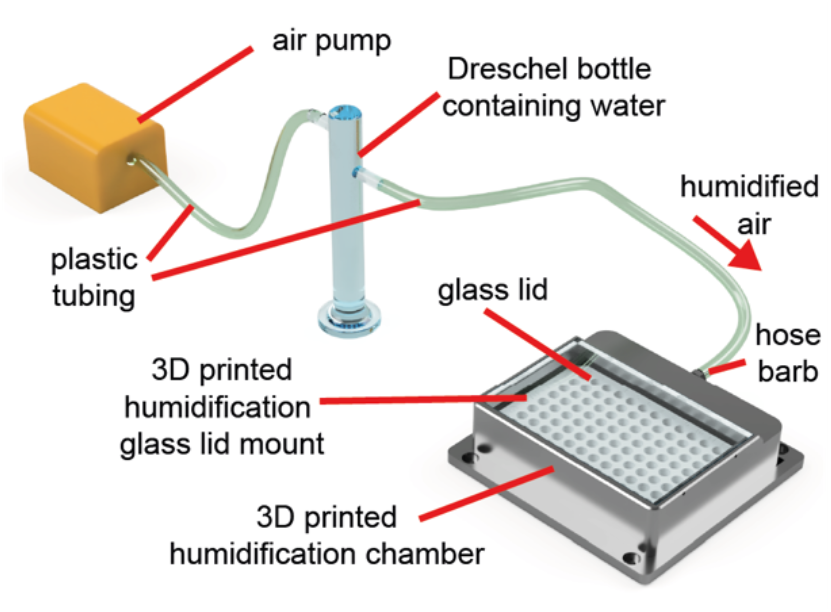
Annotated CAD render of the humidification embryo culture assembly.

**Figure 12.**
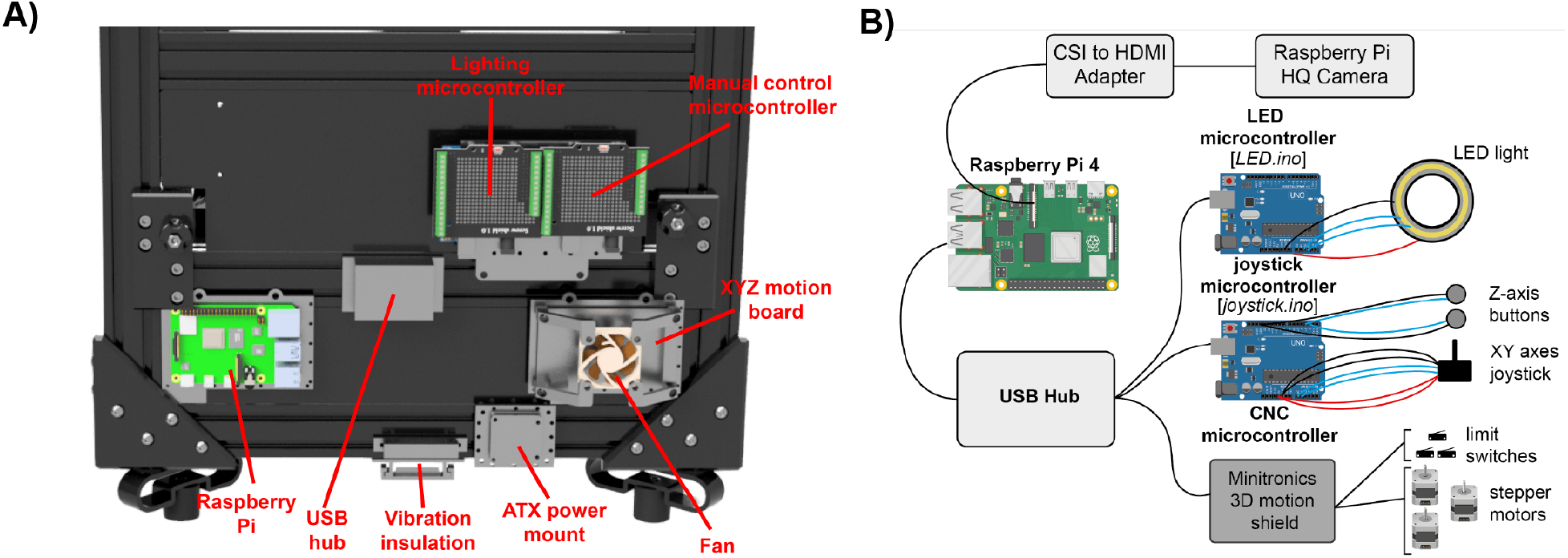
A). Annotated CAD render of the electronics mounted on the rear of the LabEmbryoCam. B) Schematic diagram of the connection between the different electronic components in the LabEmbryoCam, including the pinouts from the microcontrollers and the names of the scripts to upload to LED and joystick microcontrollers. The CNC microcontroller firmware should be uploaded following the manufacturer instructions, with the configuration files available in the project repository. A disk image is provided for flashing a microSD card with the Raspberry Pi Raspbian operating system, with all dependencies installed. Mouse, keyboard and monitor should be connected to the prior to powering on Raspberry Pi.

### Electronics

Electronics are mounted at the rear of the LabEmbryoCam, onto specifically designed 3D printed mounting parts. All cabling to the LabEmbryoCam should run through the two vibration insulation brackets on the back and bottom of the instrument - to prevent environmental vibration from impacting image acquisition (Figure 12A).

Lighting, joystick/buttons, and motor control are provided using microcontrollers, all running their own scripts which must be uploaded. See this guide for help on uploading the lighting, joystick and xyz motion scripts to microcontrollers, using the freely available Arduino IDE: https://support.arduino.cc/hc/en-us/articles/4733418441116-Upload-a-sketch-in-Arduino-IDE

#### Lighting microcontroller

Lighting is controlled *via* an Arduino UNO microcontroller, with a relatively simple script. The LuMini LED range provide a high and fairly uniform lighting pattern, at a low cost. However, there is significant scope for alternative lighting solutions.

The script for uploading to the LED microcontroller is accessible in the LED_microcontroller folder in the LabEmbryoCam software directory. Colour values can be defined within this script (default is white), and brightness is controlled via the LabEmbryoCam web application.

#### Manual input controller

The XY joystick and Z button controls connect directly to the analog and digital inputs of the manual input microcontroller (Arduino Uno) which passes commands to the Raspberry Pi via USB. The script to run on the manual input microcontroller is accessible within the LabEmbryoCam software directory in thejoystick_microcontroller folder. The wiring between the microcontroller, joystick and Z buttons are detailed in the joystick_microcontroller script.

#### XYZ motion controller

The LabEmbryoCam makes use of a CoreXY (https://en.wikipedia.org/wiki/CoreXY) style of motion control and this can be achieved using a range of different microcontroller boards from both CNC and 3D printing suppliers. The Minitronics V2 board is used here due to its low cost and integrated stepper motor drivers (reducing setup complexity). More sophisticated 3D printing, or CNC motion microcontrollers would enable higher resolution (*via* microstepping) and quieter operation but comes at greater cost.

Firmware and configuration files for the Minitronics board are accessible in the XYZ_microcontroller folder of the LabEmbryoCam software directory. The XYZ microcontroller is more sophisticated than the Arduino UNOs used for the lighting and manual input control and requires some different settings when uploading the scripts using the Arduino IDE. Further information can be found here, in the ‘Configuring Arduino’ section: https://reprap.org/wiki/Minitronics_20

#### Raspberry Pi

A Raspberry Pi 4 8GB can run the LabEmbryoCam smoothly, but it should also be possible to run on any computer on which the Raspberry Pi camera and Dash are supported-such as the NVIDIA Jetson single board computers. On other systems, the web application could be used, if camera drivers and accompanying camera web application camera modules were created. A heatsink and fan cooling for the Raspberry Pi are suggested for optimal performance, and a powered USB hub is required to prevent excessive power draw from the USB ports leading to unreliable hardware connections. The camera should be connected to the Raspberry Pi using an HDMI-CSI adapter (details in bill of materials) - to enable sufficient distance from the camera to the Raspberry Pi, mounted on the rear of the instrument.

Follow manufacturer instructions for mounting both the heatsink and CSI-HDMI adapter to the Raspberry Pi.

The Raspberry Pi can be flashed with the operating system image in the data repository, containing all dependencies installed. This can then be loaded onto a microSD card using the Raspberry Pi Imager software - https://www.raspberrypi.com/software.

The lighting, manual input, and XYZ microcontrollers should all be attached to the Raspberry Pi, *via* the USB hub, which is itself then connected to a USB3 port of the Raspberry Pi. The Raspberry Pi should have a mouse, keyboard and a monitor attached. Note that the Raspberry Pi has a microHDMI, so an adapter is required to attach to a standard HDMI cable (included in the bill of materials is attached).

The Raspberry Pi is powered by the 5 V power supply from the ATX splitter, connected to a USB-C cable. This can be made by simply cutting a USB-C cable, stripping the 5V and earth cables and connecting these to the ATX power supply 5V outputs.

### Design considerations

Reduced cost, scalability and reliability were central considerations during the design of the LabEmbryoCam. Inspiration is taken from CoreXY style 3D printers for the style of motion and the choice of readily available components. The instrument presented here uses low-cost stepper motors and stepper driver boards, and relies primarily on 3D printed brackets, however these are easily modifiable design considerations. Silent stepper drivers could be used to reduce the noise, and to increase the precision via greater micro stepping. The frame could be built using blind joints for attaching the aluminum extrusion together. Furthermore, the size of the instrument was configured to enable scanning of multiwell plate formats. However, the LabEmbryoCam size could easily be scaled both up or down, by using different length linear rails and lengths of aluminium extrusion.

### Safety Concerns

The LabEmbryoCam provides X, Y and Z movement and there is therefore the danger of entrapment of hands. This is limited by the use of aluminium extrusion to surround the instrument, with the further potential for mounting acrylic panels. The use of belts for X and Y movement also results in stalling of the motion if significant resistance is encountered, such as is the case if an obstacle is met.

Electronics in the instrument are powered at 5V, with the exception of the XYZ stepper driver board, which requires a 12V input. An ATX power supply splitter provides both a 5V and 12V output. The USB hub should be powered via its own manufacturer provided power supply. Both the Raspberry Pi and XYZ stepper driver board should be fitted with heatsinks, and actively cooled with fans. It should be noted that the Raspberry Pi’s performance will likely be reduced if not cooled.

Care should be taken if the LabEmbryoCam is used in particularly wet environments, and consideration should be given to enclosing the electronics on the back of the instrument.

## 6. Operation instructions Setting LabEmbryoCam up

- Plug the LabEmbryoCam into the mains power supply. If configured as detailed here, the LabEmbryoCam requires two power sources – a 12V connection to the ATX splitter (provided *via* the power brick included with the ATX splitter), and a barrel connection to the USB hub (provided with the USB hub).
- Follow the software installation instructions above.
- Attach an HDMI, keyboard and mouse to the LabEmbryoCam. If possible, run these cables through the vibration dampener at the rear of the instrument. For optimal performance, image acquisitions should be saved somewhere other than the microSD card, such as a 2.5” SSD. A cable adapter for a 2.5” SSD is included in the bill of materials, and a 3D printed bracket is provided for mounting this securely on the rear of the instrument (Figure 12).
- If an internet connection is available, either *via* ethernet, or wifi, the LabEmbryoCam can send notifications of the progress of an acquisition – enabling remote monitoring. Wifi can be setup using the Raspbian operating system. Some organizational networks are not compatible with Raspberry Pi.

### Launching user interface

- Navigate to the folder containing the LabEmbryoCam program and select ‘Open in Terminal’. Once the Terminal window is open, type: python app.py
- Once you have started the LabEmbryoCam user interface you will be given an address to open in your browser and once opened you should see the user interface (Figure 13A).
- We can now proceed with getting the hardware set up for running an experiment. The first thing we will do is to home the XYZ stage. This is essential to ensure that the correct origin is used when finding and creating positions. To do this, you must click the XYZ tab to switch from the camera to the XYZ window, and then click on the ‘Home’ button. Before homing the stage, make sure there are no objects that could obstruct the movement of the stage. Also, make sure not to use the app whilst the stage is homing as this could interfere with the process.

**Figure 13.**
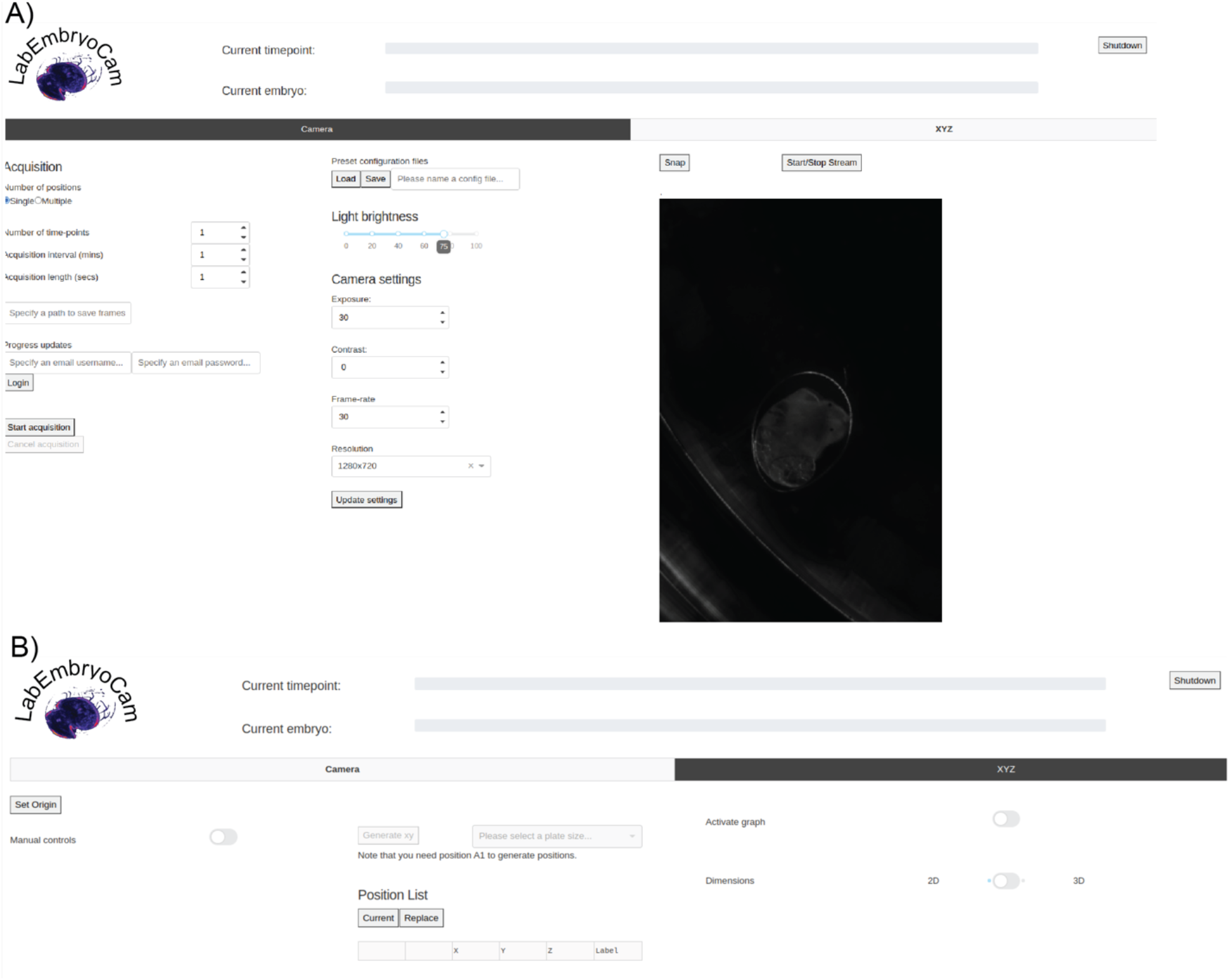
A) Camera panel and B) XYZ panel of the LabEmbryoCam web application.

### Setting up an experiment

- Once the stage is homed (origin set), you can now make use of the manual controls to move the stage to desired positions. To enable the joystick and Z buttons, toggle the ‘Manual Controls’ switch in the XYZ tab of the UI (Figure 13B)
- In order to record positions for an experiment, you need to first start a camera live stream to ensure our positions are correct. This can be done by switching back to the Camera tab and clicking the Start/Stop Stream button.
- Now that you have a live stream from the camera, we can position the stage over a desired position by using the manual controls and the live stream as feedback. You must first activate the ‘Manual Controls’ toggle on the XYZ page to enable the physical controls on the front of the LabEmbryoCam.
- Before changing any of the camera settings you must exit the live stream so that your changes can take effect. When you have finished choosing your desired settings, press the Update Settings button before starting the live stream again.
- Once you have a position that you want, click the ‘Add’ button in the XYZ tab. Note that you can either record your own positions, or save time by making use of the automated XYZ multiwell plate position generator.
  o To automatically generate positions for well of a multiwell plate, first move to the centre of well ‘A1’ and click ‘Current’ – give this well the label ‘A1’. You can then choose a multiwell plate format from the dropdown at the top of the XYZ tab – and click ‘Generate xy’.
  o If not all of the wells that are generated are required, you can delete these rows from the position list.
  o You can also modify the positions using the manual controls, and ‘Replace’ button.
- To more easily visualize and navigate a multiwell plate, click the ‘Activate Graph’ toggle. This will present you an interactive map enabling you to move between positions by clicking twice on any point.
- Once all desired positons are listed in the XYZ position table, the next step is to enter the parameters for the acquisition before starting the experiment.
- Switch to the camera tab, and adjust the following parameters:
  o Number of positions - Whether you would like to capture footage for only the current position (‘Single’) or all the positions you have recorded (‘Multiple’).
  o Number of timepoints - How many acquisition iterations you would like the system to complete. An iteration consists of capturing footage for all of the specified positions.
  o Acquisition interval - How long to wait between each timepoint in minutes.
  o Acquisition length - How long to capture video for each position, at each timepoint.
  o Save folder - The full file path to the directory where you would like to save video.
  o Progress updates - Username and password for gmail specific to your unit. Click the login button to send progress updates to this email account after every timepoint.
  o After entering the parameters for your experiment, the last step is to press the Start Acquisition button to begin the experiment. Note that you can cancel an acquisition in progress if you would like to make adjustments to the parameters used by the software.

### Output of an experiment

- Individual AVI videos are saved for each time point within a folder labelled with the XYZ position label. Metadata on the acquisition and timings are also stored in capture_data.csv and time_data.csv (https://doi.org/10.5281/zenodo.7575249).
- Videos for each embryo can be compiled into a longer video covering the duration of the experiment using the script ‘compile_videos.py’ located in the analysis scripts folder of the LabEmbryoCam software.
- Either raw video from the LabEmbryoCam, or the compiled videos can be played using the ImageJ variant Fiji, or VLC media player. In the analysis_scripts folder within the LabEmbryoCam directory, there is a simple user interface for playing either individual videos via ‘timepoint_viewer.py’, or multiple timepoints via multi-timepoint_viewer.py, for a particular embryo.
- The video from the LabEmbryoCam can be used in a number of downstream analytical processes described in previously published research, including optical flow [5], heart rate detection [9], size, shape and movement measurement [7], or energy proxy traits [11].

## 7. Validation and characterization

### Experiment setup and testing

LabEmbryoCam was run for 48 h recording the development of nine early hippo stage embryos of the freshwater gastropod Radix balthica, generating forty eight 20 second videos, demonstrating the capability of the instrument to run unsupervised for extended periods (https://doi.org/10.5281/zenodo.7575249).

To assess the capability of video produced using the LabEmbryoCam for automated image analysis, it was analysed using the open-source Python package EmbryoCV [7]. EmbryoCV produced effective measures of change to size, movement, and also energy proxy traits – energy in the spectrum of fluctuating pixel values, an emerging transferrable phenotyping approach [8, 11] (Figure 14). The LabEmbryoCam was effective in generating video of a quality that enabled the extraction of continuous individual-level physiological data, including growth, movement and energy proxy traits.

**Figure 14.**
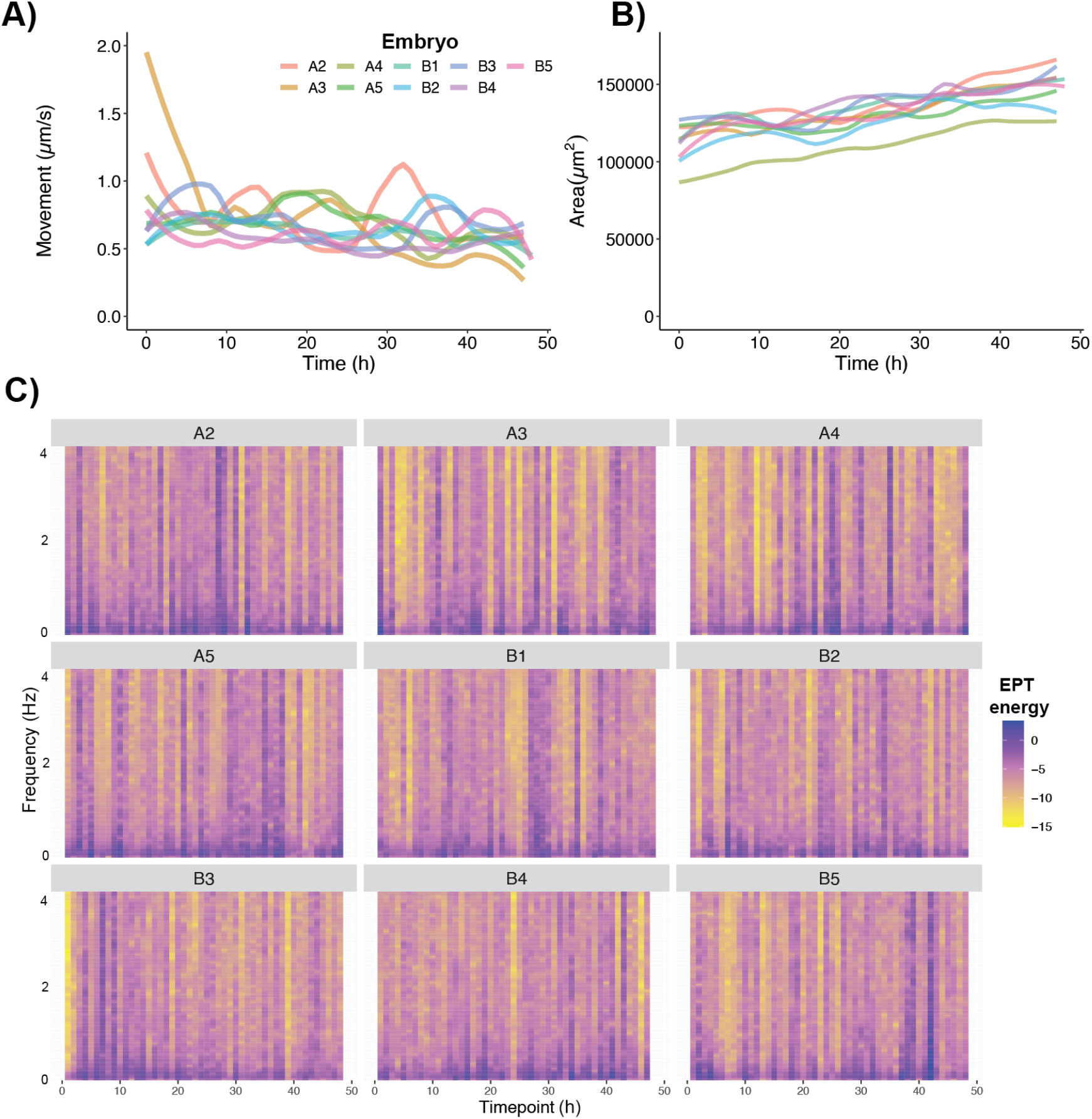
A) Developmental change in speed of the movement of individual embryos. B) Growth of individual embryos. C) Levels of energy in the EPT spectra of individual embryos during the course of the experiment.

### Humidification

The humidification chamber is an optional addition to the LabEmbryoCam to overcome the common evaporation issue associated with multiwell plates. Humidity > 90 % was achieved in the humidified airspace above a 96 multiwell plate (DHT22 humidity sensor) using a flow rate of 1 L/min for the duration of a 48 h test.

### Camera and lens performance

The performance of the camera was assessed by examining the metadata from the 48 h camera to investigate consistency of frame to frame timings (Figure 14A) and video duration (Figure 14B). The camera was effective at operating at a range of resolutions, and corresponding upper frame rates (Table 2), and presets for these are included in the LabEmbryoCam webserver.

**Table 1.** Frame rates for acquisition of video from the Raspberry Pi HQ camera, connected to a Raspberry Pi 4 via HDMI-CSI adapter, controlled using the LabEmbyoCam webserver.

| Resolution (pixels) | Lower frame rate(fps)<br>> 10 ms exposure | Upper frame rate (fps)<br>< 10 ms exposure |
| --- | --- | --- |
| 640 x 480 | 30 | 90 |
| 1280 x 720 | 30 | 90 |
| 1920 x 1080 | 30 | 40 |
| 256 x 256 | 30 | 90 |
| 512 x 512 | 30 | 90 |
| 1024 x 1024 | 30 | 80 |
| 2048 x 2048 | 30 | 20 |

### XYZ movement accuracy

With the aim of testing the accuracy of camera movements, two tests were carried out. We selected two positions to take 100, 1 second videos per position every 5 minutes. A distance of 1mm between positions was used for the first test and a distance of 100mm used for the second (Figure 14C). To assess the accuracy of camera movements, images were taken of a calibration slide before and after movement. Using the open source image analysis software ImageJ [12], distances moved (μm) were calculated and used to assess the accuracy of movement relative to the known distances.

**Figure 14.**
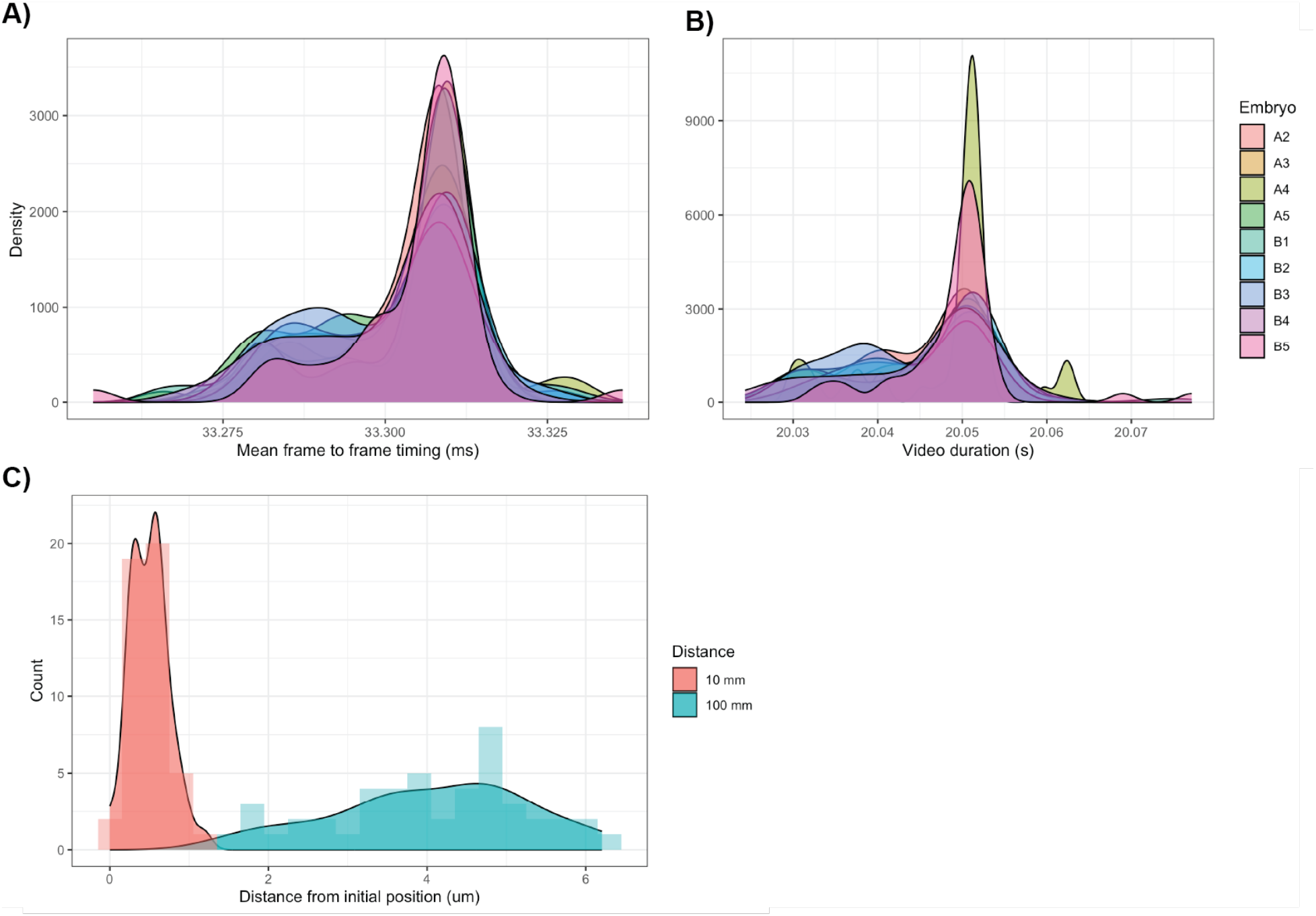
A) Density of mean frame to frame timings and B) video duration for nine embryos, imaged for 20 seconds at 30 fps, hourly during a 48 h period. C) Spatial repeatability of X and Y movements during two tests of 48 movements of 10 mm and 100 mm of the LabEmbryoCam in the X and Y axis directions.

## CRediT author statement

**Ziad Ibbini:** Software, Conceptualization, Methodology, Writing **Maria Bruning:** Methodology, Writing **Sakina Allili:** Methodology **Luke Holmes:** Methodology, Validation, Writing **Ellen Tully:** Conceptualization, Writing **Jamie McCoy:** Methodology, Writing **John I. Spicer:** Conceptualization, Writing **Oliver Tills:** Conceptualization, Methodology, Writing, Project administration

## Acknowledgments

This work was funded by a NERC ARIES funded PhD studentship awarded to James McCoy, a University of Plymouth funded PhD studentship awarded to Ziad Ibbini, a Santander Undergraduate Exchange Scholarship awarded to Maria Bruning and a UKRI Future Leaders Fellowship (MR/T01962X/1) awarded to Dr Oliver Tills.

## References

1. D. Houle, D.R. Govindaraju, S. Omholtm, Phenomics: the next challenge. Nature Reviews Genetics 11, 855–866 (2010).

2. D. Kültz, Evolution of cellular stress response mechanisms. Journal of Experimental Zoology Part A: Ecological and Integrative Physiology 15, 217 (2020).

3. R.T. Furbank, M. Tester, Phenomics--technologies to relieve the phenotyping bottleneck. Trends in Plant Science 16, 635–644 (2011).

4. L.L.M. Pandori, C.J.B. Sorte, The weakest link: sensitivity to climate extremes across life stages of marine invertebrates. Oikos 128, 621–629 (2018).

5. O. Tills, T. Bitterli, P. Culverhouse, J.I. Spicer, S.D. Rundle, A novel application of motion analysis for detecting stress responses in embryos at different stages of development. BMC Bioinformatics 14, 37 (2013).

6. O. Tills, S.D. Rundle, J.I. Spicer, Parent-offspring similarity in the timing of developmental events: an origin of heterochrony? Proceedings of the Royal Society: Biological Sciences, 280, 20131479–20131479 (2013).

7. O. Tills, J.I. Spicer, A. Grimmer, S. Marini, V.W. Jie, A high-throughput and open-source platform for embryo phenomics. PLoS Biology 16, e3000074 (2018).

8. O. Tills, J.I. Spicer, Z. Ibbini, S.D. Rundle, Spectral phenotyping of embryonic development reveals integrative thermodynamic responses. BMC Bioinformatics 22, 232 (2021).

9. Z. Ibbini, J.I. Spicer, M. Truebano, J. Bishop, O. Tills, HeartCV: a tool for transferrable, automated measurement of heart rate and heart rate variability in transparent animals. Journal of Experimental Biology (2022) doi:10.1242/jeb.244729.

10. C. Dwane, S.D. Rundle, O. Tills, E. Rezende, J. Galindo, E. Rolán-Alvarez, M. Truebano, Divergence in Thermal Physiology Could Contribute to Vertical Segregation in Intertidal Ecotypes of Littorina saxatilis. Physiological and Biochemical Zoology 94, 353–365 (2021).

11. O. Tills, L.A. Holmes, E. Quinn, T. Everett, M. Truebano, J.I. Spicer, Phenomics enables measurement of complex responses of developing animals to global environmental drivers. Science of the Total Environment 159555 (2022).

12. J. Schindelin, I. Arganda-Carreras, E. Frise, V. Kaynig, M. Longair, T. Pietzsch, S. Preibisch, C. Rueden, S. Saalfeld, B. Schmid, J. Tinevez, D.J. White, V. Hartenstein, K. Eliceiri, P. Tomancak, A. Cardona. Fiji: an open-source platform for biological-image analysis. Nature Methods 9, 676–682 (2012).

